# Ecological models predict narrow potential distribution for *Trioza erytreae*, vector of the citrus greening disease

**DOI:** 10.1101/2022.07.07.496964

**Authors:** Martin Godefroid

## Abstract

The African citrus psyllid, *Trioza erytreae* (Hemiptera: Triozidae), is a vector of citrus greening disease (Huanglonbing - HLB) caused by the bacterium *Candidatus liberibacter.* Native from Africa, *T. erytreae* was detected in northwestern Spain in 2014, and since then it has established along Atlantic coastal areas of the Iberian Peninsula. Given the severe bio-economic impact of HLB, an accurate assessment of the risk of potential spread of African citrus psyllid to citrus-growing regions of the Mediterranean area and the rest of the world, is urgently needed to design effective control strategies and anticipate economic losses. Therefore, I calibrated species distribution models to understand the bioclimatic characteristics that shape the distribution of *T. erytreae* and to assess the climatic suitability of the world’s major citrus-growing regions for this species under current and future climate conditions. The models identify mild summer and winter temperatures and high levels of precipitation as optimal conditions for long-term psyllid establishment. It is noteworthy that the models trained without the available occurrences in continental Europe, predict only the Atlantic coastal regions of the Iberian Peninsula as highly climatically suitable in Europe, which corresponds perfectly with the area currently invaded by the psyllid. This striking predictive accuracy lends great credibility to the model predictions. Most economically important citrus production areas in the world are predicted to be of low or moderate climatic suitability for *T. erytreae.* This research is crucial for assessing the global risk of HLB and is particularly timely for Europe where the African citrus psyllid has recently been detected.

## Introduction

Citrus greening disease (i.e. Huanglongbing in Chinese - HLB), induced by the bacterium *Candidatus liberibacter,* is a major problem for citrus cultivation (Bové 2006). Common symptoms associated with HLB include leaf yellowing, defoliation, decreased root abundance, twig dieback, production of small, irregularly shaped and bitter fruits, and a general decline health, which eventually leads to plant death. Since there is no treatment available to cure HLB-infected trees, the economic losses induced by the disease are severe for the citrus industry. The economic costs are related to substantial decreases in production as well as the use of plant protection products and the implementation of costly control strategies (Hodges and Spreen 2012; Spreen et al. 2014).

The pathogen *C. liberibacter* is mainly transmitted from one plant to another by insects. Two main vectors of HLB are currently recognized, namely the Asian citrus psyllid *Diaphorina citri* Kuwayama (Hemiptera: Liviidae) and the African citrus psyllid *Trioza erytreae* Del Guercio (Hemiptera: Triozidae). The Asian citrus psyllid is native to Asia and has spread to most of economically-important citrus-growing regions of the world - including North America, the Caribbean, the Middle East, and South America - where it is responsible for severe HLB epidemics (CABI 2021). The African citrus psyllid is native to Africa and has spread to the Indian Ocean islands, the Canary Islands, the island of Madeira, as well as to continental Europe (CABI 2021). However, *T. erytreae* is still absent from the main citrus-growing regions located in Americas, Asia, and Australia. In continental Europe, the African citrus psyllid was first officially detected in 2014 in the northwestern parts of the Iberian Peninsula (Otero et al. 2015). Since then, the insect has remained confined to the Atlantic coastal regions of northern Spain and Portugal and has not spread to economically important citrus-growing regions of the Mediterranean basin (Arenas-Arenas et al. 2019).

The African citrus psyllid feeds mainly on plants belonging to the Rutaceae family (Moran 1968). The species does not diapause and can reach a total of 6-8 generations when environmental conditions are optimal (Catling 1972; Tamesse and Messi 2004). However, under suboptimal conditions, the annual number of generations decreases sharply (Tamesse and Messi 2004). Depending on temperature conditions, eggs hatch in 1 to 2 weeks while 2 to 8 weeks are required to complete the five nymphal stages. Eggs and nymphs are thought to be the most temperature- and humidity-sensitive life stages; in particular, populations normally experience high mortality of eggs and early nymphal stages when conditions are hot and dry (der Merwe 1923; Moran VC & Blowers 1967; Catling 1969; Green and Catling 1971; Samways 1987). Adults can live for several months and are able to cope with adverse conditions of drought and heat (Samways 1987).

Risk assessment of a vector-borne disease ideally involves independently predicting the environmental tolerances of the pathogen and its vectors (Godefroid et al. 2021; Gimenez-Romero et al. 2022). Given the economic importance of HLB, there is an urgent need to assess the potential risk of *T. erytreae* spread into citrus-growing regions in Europe and the rest of the world. Therefore, the objective of this study is to understand the climatic factors that determine the spatial distribution of the African citrus psyllid and to predict the climatic suitability of the world’s major citrus growing regions for this species. To achieve this goal, I collected detailed data on the distribution of *T. erytreae* and calibrated bioclimatic species distribution models (SDMs; Peterson et al. 2011) to describe the environmental tolerances of this species and assess climatic suitability in Europe as well as in the rest of the world’s important citrus producing regions.

## Material and Methods

### Occurrence data

I extracted presence data from the scientific literature and the freely available GBIF (Global Biodiversity Information Facility database (Gbif.org 2021). I also obtained occurrence records from online reports available on the websites of the Province of Galicia (Xunta de Galicia 2020) and the Portuguese Directorate General of Food and Veterinary Medicine (Direção-Geral de Alimentação e Veterinária 2021). I checked the reliability of each record and discarded unreliable occurrences. A total of 2,771 presence records were obtained (Fig. 1). After removing duplicate records (i.e., allowing only one presence record in each pixel of the bioclimatic rasters used for modeling), the final dataset included 1,500 occurrences.

**Figure 1.**
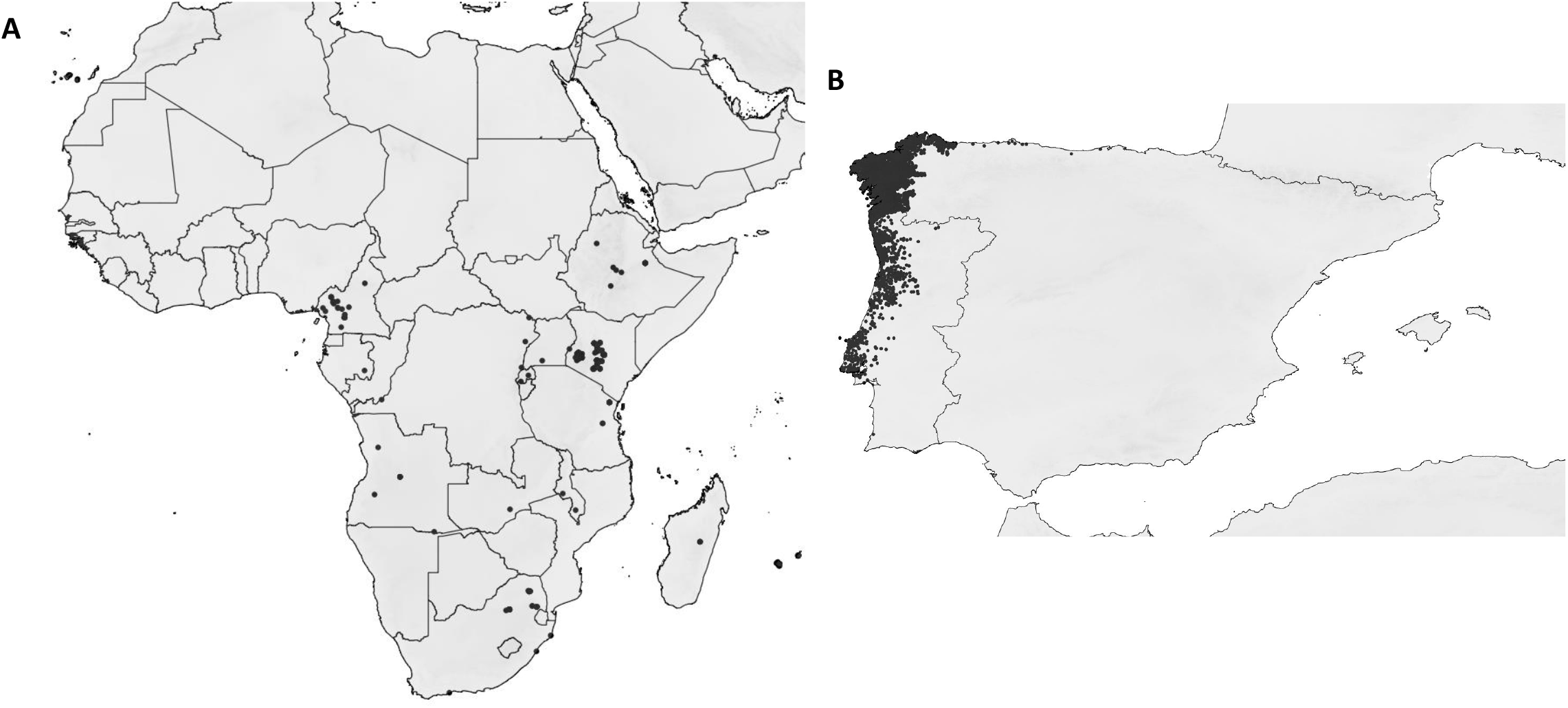
Presence records for the African citrus psyllid *Trioza erytreae* in (A) Africa, Indian Ocean Islands, Atlantic Ocean Islands and (B) mainland Europe. Presences of *T. erytreae* are represented by black dots.

### Bioclimatic data

I extracted global bioclimatic rasters from the CHELSA database, which reflects historical climate trends for the period 1979-2013 (Karger et al. 2017). The rasters were downscaled to a 2.5 arc minutes resolution. Four bioclimatic descriptors thought to reflect putative climate stress for this species were used in the modeling framework, namely the mean temperature of the warmest quarter of the year (bio10), the mean temperature of the coldest quarter of the year (bio11), the precipitation of the warmest quarter of the year (bio18), and the precipitation of the coldest quarter of the year (bio19). I ensured that these four covariates were not highly correlated in the calibration area (i.e., a Pearson correlation index < 0.7).

To predict the impact of climate change on the potential distribution of *T. erytreae,* I used estimates of future climate conditions for the period 2040-2060 simulated by two global climate models provided by the Intergovernmental Panel on Climate Change Fifth Assessment Report, namely the Model for Interdisciplinary Climate Research version 5 MIROC5 and the Canadian Earth System Model version 2 (CanESM2) (Watanabe et al. 2011; Swart et al. 2019). I used a moderate scenario of future greenhouse gas emissions (rcp45 scenario).

### Calibration and evaluation of models

I used the algorithm Maxent to model the environmental tolerances of the African citrus psyllid (Phillips et al. 2006). This algorithm is a widely used presence-only approach that ranks among the best performing SDM techniques. (Wisz et al. 2008). I allowed only linear and quadratic features to ensure easy interpretation of the models and to avoid fitting overly complex species-covariate relationships and overparameterizing the models (Merow et al. 2013). I calibrated the models using only available occurrences in Africa, Indian Ocean islands, and Atlantic Ocean islands. I generated 20,000 background points for model calibration (i.e., localities with unknown species presence status) generated in Africa, the Middle East, Indian Ocean islands, and Atlantic Ocean islands (Appendix S1). Models were fitted using a subset of 90% of these data (calibration dataset) and evaluated using the remaining 10% of data (evaluation dataset). I evaluated the models by calculating the area under the receiving operator characteristic curve (AUC) (Fielding and Bell 1997) and the True Skill Statistics (TSS) (Allouche et al. 2006). The models were also evaluated using available occurrence records in continental Europe and 1,000 background points generated in the Iberian Peninsula, which can be considered as a spatially independent evaluation procedure. The models were replicated 5 times and the average climatic suitability between model replicates was mapped. The importance of each variable in models was estimated following the approach proposed by Thuiller et al (Thuiller et al. 2009). Models were fitted and evaluated using the *biomod2* R package (Thuiller et al. 2016).

## Results

The models produced excellent evaluation measures (mean AUC = 0.89 ± 0.03; mean TSS = 0.69 ± 0.09) when evaluated with the evaluation dataset available in the calibration area. The models also provided excellent evaluation metrics when evaluated with the independent evaluation dataset available in continental Europe (mean AUC = 0.97 ± 0.002; mean TSS = 0.6 ± 0.19). Bioclimatic models identified summer temperature (bio10) as the most important covariate in explaining the distribution data (mean variable importance = 0.77 ± 0.03), followed by winter temperature (bio11; mean variable importance = 0.16 ± 0.01) and precipitation in the warmest quarter of the year (bio18; mean variable importance = 0.06 ± 0.01). Precipitation in the coldest quarter of the year (bio19) made a negligible relative contribution to the model (mean variable importance = 0.006 ± 0.004). The models modeled unimodal responses of African citrus psyllid to winter and summer temperatures (Appendix S2). The models modeled a positive linear-like effect of increased rainfall during the warmest and coldest quarters of the year on the probability of *T. erytreae* occurrence (Appendix S2).

Bioclimatic models predicted that a very small part of Europe is currently highly climatically suitable, which ideally matches the invaded range of *T. erytreae* in the Iberian Peninsula (Fig. 2). In Europe, the regions predicted to be most suitable encompass the cool, humid areas of the northern parts of the Iberian Peninsula, i.e. the Atlantic coastal regions of Portugal and Spain (Figs. 2 & 3). The dry and hot regions of the Mediterranean basin, where economically important citrus cultivation is generally practiced (e.g., lowlands of southern and eastern Spain, lowlands of southern Italy and Greece) were considered moderately or poorly suited (Figs. 2 & 3). Bioclimatic models did not predict a significant increase in climatic suitability for *T. erytreae* in these European citrus-growing regions for the period 2040-2060 (Fig. 4 & Appendix S3).

**Figure 2.**
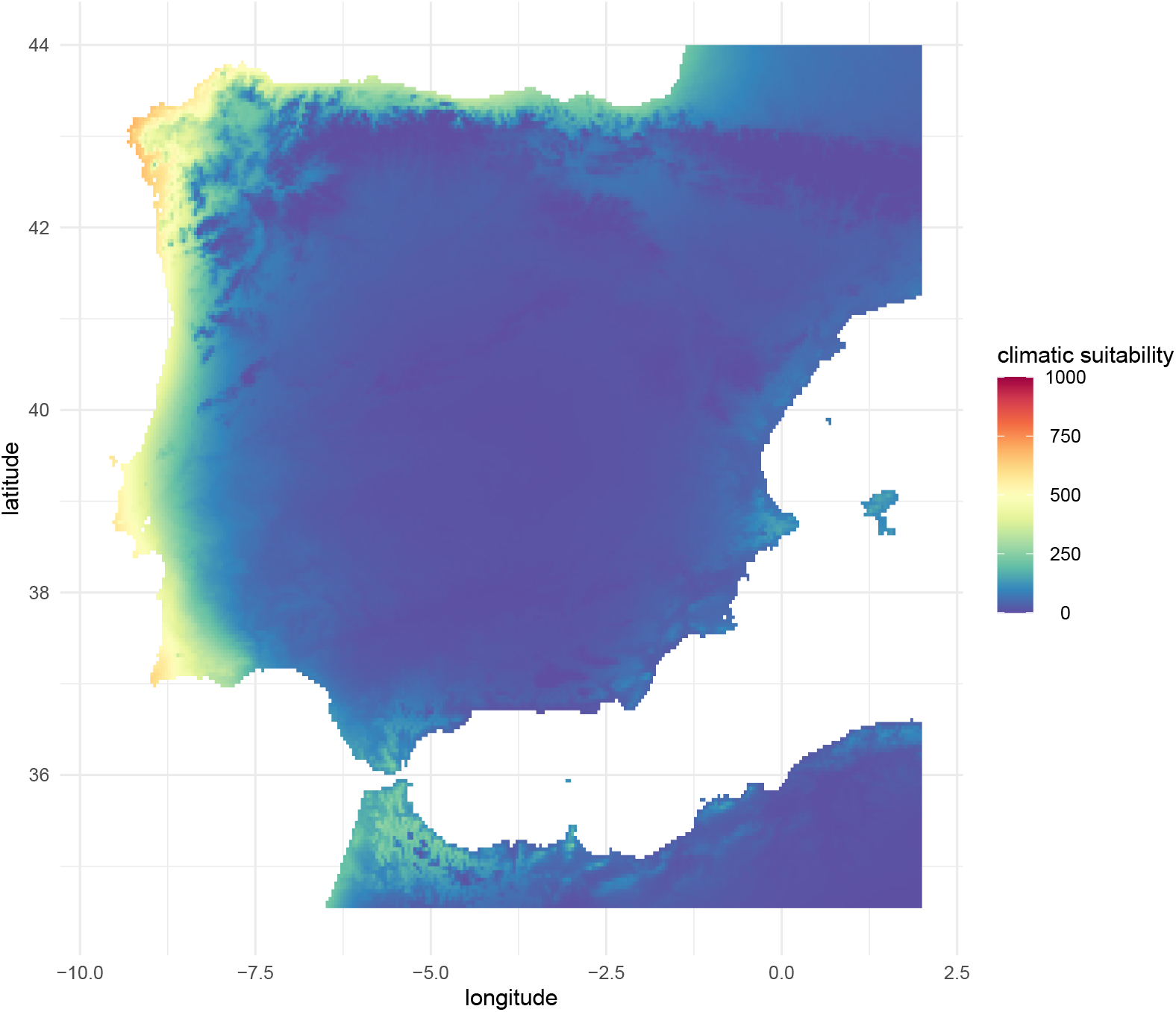
Maxent-derived predictions of current climate suitability in the western Iberian Peninsula for the African citrus psyllid *Trioza erytreae.* Maps represent the average predictions among 5 replicates.

Major economically important citrus-growing areas in the rest of the world (FAO, 2019) were globally predicted to be poorly suitable, e.g. Florida and California (USA), lowlands in southeastern China, India, lowlands in Mexico, lowlands in tropical Brazil, and coastal regions in southern Turkey or Egypt (Fig. 3). Bioclimatic models did not predict a significant increase in climate suitability for *T. erytreae* in these regions for the period 2040-2060 (Fig. 4 & Appendix S3).

**Figure 3.**
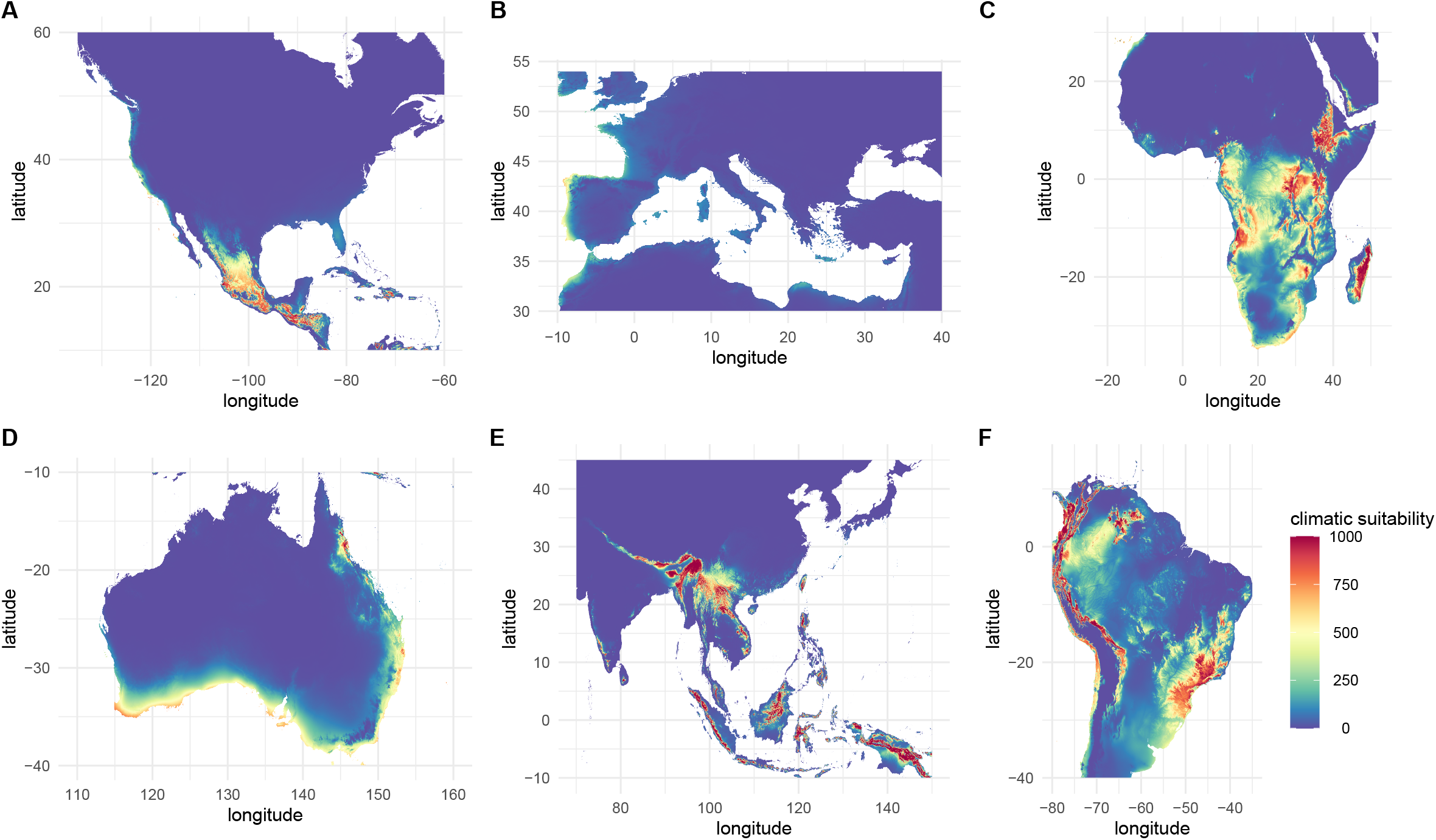
Maxent-derived predictions of current climate suitability of some of the major citrus-growing regions of the world for the African citrus psyllid *Trioza erytreae* (A: North America; B: Europe and North Africa; C: South Africa; D: Australia; E: Asia; F: South America). Maps represent the average predictions among 5 model replicates.

**Figure 4.**
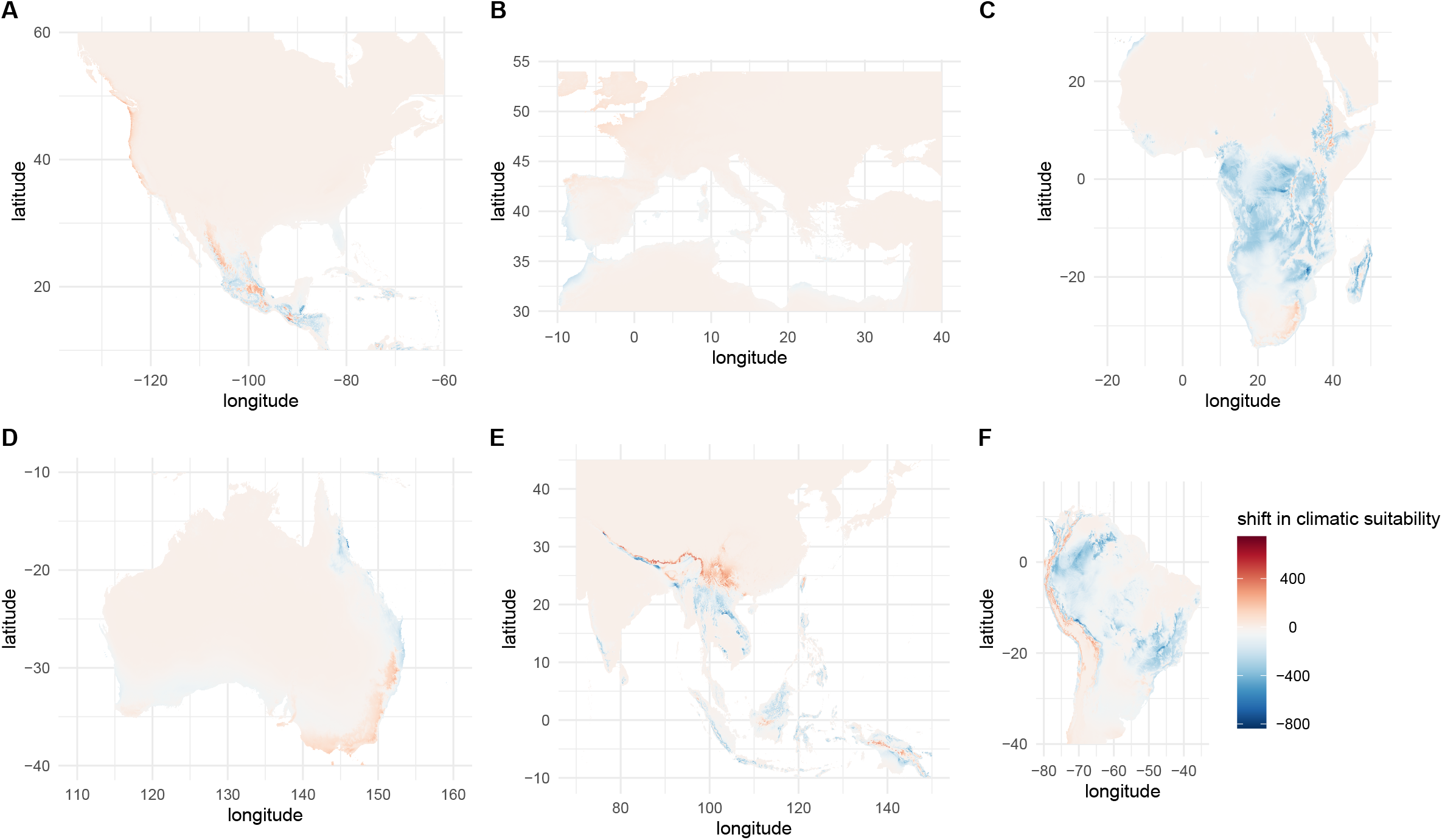
Maxent-derived predictions of increase/decrease in climate suitability for the African citrus psyllid *Trioza erytreae* by the period 2040-2060 under the rcp45 scenario of future climate conditions (MIROC5 global circulation model). Maps represent the subtraction between the predicted current climate suitability and that simulated under future climate conditions.

## Discussion

### Reliability of models’ outputs

The Maxent-derived modeled response curves to bioclimatic covariates are consistent with currently available knowledge of the physiology of *T. erytreae.* Indeed, the Maxent models identified a negative effect of high temperatures on the probability of African citrus psyllid occurrence, which is consistent with early field observations (Moran VC & Blowers 1967; Green and Catling 1971; Samways 1987) and recent laboratory-derived evidence that this species is heat-sensitive (Aidoo et al. 2022). Similarly, models have identified a negative correlation between low winter temperatures and the likelihood of African citrus psyllid occurrence, consistent with the apparent lack of cold-induced diapause in this psyllid and the long-standing assertion that a temperature of 10 degrees Celsius is the lower threshold for development of various life stages of this species (Catling 1969, 1973). Similarly, no complete development of *T. erytreae* was observed in climatic chambers kept constantly at 10 degrees Celsius (Aidoo et al. 2022). Bioclimatic models also suggest that high amounts of rainfall during both the warmest and coldest periods of the year favor the presence of *T. erytreae,* confirming that dry conditions are not optimal for the multiplication of this species (Moran VC & Blowers 1967; Green and Catling 1971; Samways 1987).

The estimates of climatic suitability presented in the herein study are consistent with the known distribution of the African citrus psyllid outside of continental Europe, i.e., this species is abundant in cool areas and absent or at least rare in the warm lowlands of sub-Saharan Africa (Aubert et al. 1988; Tamesse et al. 1999; Aidoo et al. 2019) and La Réunion Island (Aubert et al. 1980). Strikingly, the models predicted the area invaded by *T. erytreae* in continental Europe with high accuracy (Fig. 2). I recall that the distribution data from mainland Europe were not considered in model calibration and were only used as an independent evaluation dataset. Such accuracy in predicting a spatially independent evaluation dataset lends great credibility to the SDM-derived climate suitability estimations presented herein (Randin et al. 2006). According to models, the current spatial pattern of spread of *T. erytreae* along the Iberian Atlantic coast likely reflects the unique climatic conditions encountered in these regions i.e., mild temperatures in both summer and winter (Godefroid et al. 2016), which ideally fit the environmental tolerances of the African citrus psyllid.

To my knowledge, the present study represents the first attempt to provide a world map of the potential current and future climatic suitability for the African citrus psyllid by fitting correlative bioclimatic SDMs based on the occurrence data available in its native range. For the Iberian Peninsula, my models provide different results from two recently published SDM-derived predictions (Benhadi-Marín et al. 2020, 2022). I argue that the present study provides a more reliable SDM-derived estimate of the climate niche of *T. erytreae.* On the one hand, the study published by Benhadi-Marín et al. (2020) presents two major methodological caveats that might explain why their SDM-derived climatic suitability estimates do not ideally match the currently observable spread pattern of the psyllid in Portugal and Spain. Firstly, calibration and evaluation of their models were both only based on the distribution data available in the recently invaded Iberian Peninsula, which can cause a risky loss of information (Elith et al. 2010). Second, their models were calibrated with a single explanatory climatic covariate (i.e. precipitation in the coldest quarter of the year) and therefore do not consider the thermal tolerances of the psyllid, which are however crucial to explain its geographical range. Finally, precise information on the spatial extent where background points were generated in Maxent fitting is not provided by Benhadi-Marín et al. (2020), which makes reliability of their models hard to interpret. On the other hand, the study published by Benhadi-Marín et al. (2022) has a major caveat, namely that the authors used a SDM-derived estimate of the climatic suitability of *Citrus* spp. as a surrogate for the potential distribution of *T. erytreae* in the Iberian Peninsula. As pointed out in this present study and published references herein, *T. erytreae* and *Citrus* spp. do not have equivalent climatic tolerances.

I warrant that caution should be exercised in interpreting the estimates of climatic suitability presented herein, because of the correlative nature of the modelling technique (Peterson et al. 2011). Moreover, we modelled the distribution of *T. erytreae* by implementing relatively simple model features to favor transferability and interpretability of models and avoid model overfitting. I suggest that mechanistic modelling approaches could likely depict a more accurate picture of the complex and interacting climatic constraints that determine the distribution of the African citrus psyllid. I also emphasize that low predicted climatic suitability should not be interpreted as a predicted “total absence” or “no risk of local establishment.” However, I argue that these SDM-derived climate suitability maps are an interesting proxy for forecasting differences in potential population densities of the psyllid and identifying the citrus-growing regions most likely for long-term establishment of this species.

### Risk of establishment in economically important citrus-cropping regions

In Europe, four Mediterranean countries are economically important producers of citrus fruits i.e. Spain, Italy, Portugal and Greece (EFSA et al. 2019). My predictions suggest that most of the citrus-growing regions of these countries are climatically little suitable for establishment of the African citrus psyllid (Figs. 2 and 3). In coastal Portugal, which was predicted to be highly suitable for *T. erytreae,* citrus cultivation is not economically important and citrus plants are generally found in home gardens. In Portugal, most of the citrus production comes from the Algarve region located in the south of the country. According to the models, the Algarve region is considered moderately adapted to the African citrus psyllid (Fig. 2). In Spain, the economically important citrus growing areas are mainly located in the provinces of Andalusia, Murcia, and Valencia. (EFSA et al. 2019), which were predicted as little suitable for *T. erytreae* (Fig. 3). Similarly, in Italy, citrus production is limited to the southern provinces, which were also predicted to be little suitable for the African citrus psyllid (Fig. 3). According to predictions considering global change simulations, the risk of longterm establishment of *T. erytreae* in the main citrus-producing regions of Europe is expected to remain relatively low over the next few decades, which is not surprising for a species preferentially found in regions characterized by cool summer temperatures.

In the rest of the world, many economically important citrus-growing regions were predicted little suitable for establishment of the African citrus psyllid, including the major producing areas of the Mediterranean coastal regions of Turkey, the lowlands of tropical Brazil, southeastern China, India, the southern United States (especially Florida and California), and the lowlands of Mexico. (FAO, 2019). As for Europe, bioclimatic models suggest that global change will not alter this pattern in a near future (Fig. 4 & Appendix S3). We recognize that a more accurate assessment of the local risk of establishment of *T. erytreae* in citrus-growing areas would require accurate, high-resolution maps of citrus orchard distribution. However, such information on a global scale is currently lacking.

### Conclusions

Overall, this study highlights that there is a high degree of spatial variation in the potential climatic suitability for the African citrus psyllid in Europe and in the major citrus producing areas of the world. The maps provided here are crucial tools for assessing the risk of global HLB outbreaks. It should be noted that the climate suitability of the world’s major citrus producing areas is relatively low and is expected to remain so in a near future. This result has crucial implication for the design of control strategies and risk assessment of HLB, especially in Europe where *T. erytreae* has been recently detected. The introduction of the African citrus psyllid reminds us that the Asian citrus psyllid *D. citri* could be introduced into Europe as it has been in the main citrus-growing regions of the world and, therefore, probably constitutes the greatest threat to citrus cultivation in Europe and most countries of the world.

## Authors’ contributions

MG designed the study, collected the data, performed statistical analyses, and wrote the manuscript.

## Data availability statement

Data are available upon request to the author

## Supplementary Information

**Appendix S1** Distribution of background points generated to model the distribution of the African citrus psyllid.

**Appendix S2** Maxent-derived modelled response curves for the four climate covariates used in model calibration.

**Appendix S3** Maxent-derived predictions of increase/decrease in climate suitability for the African citrus psyllid *Trioza erytreae* by the period 2040-2060 under the rcp45 scenario of future climate conditions (canESM2 global circulation model). Maps represent the subtraction between the predicted current climate suitability and that simulated under future climate conditions.

