## Supplementary figures and images for "Ecological models predict narrow potential distribution for *Trioza erytreae*, vector of the citrus greening disease"

### Appendix S1

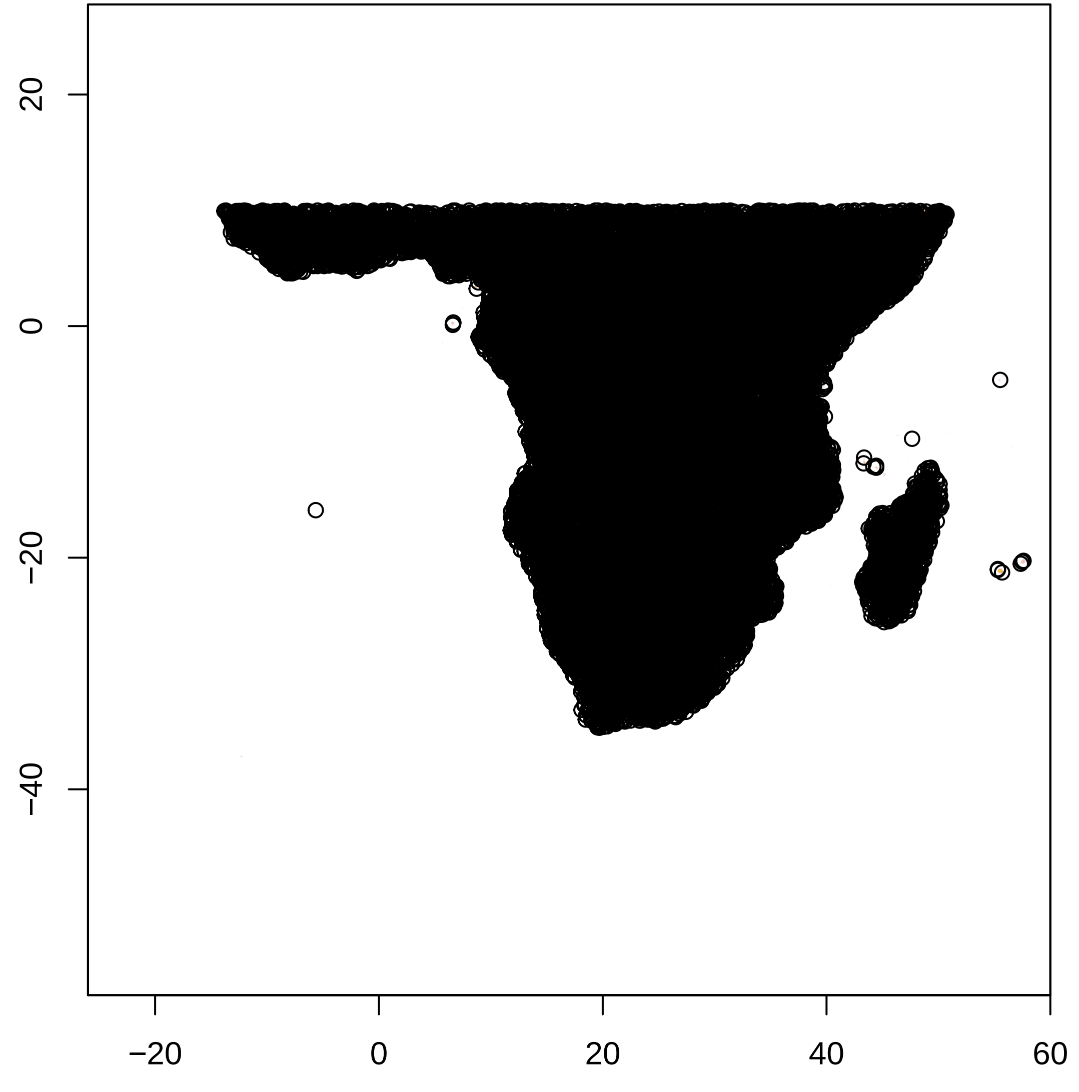

### Appendix S2

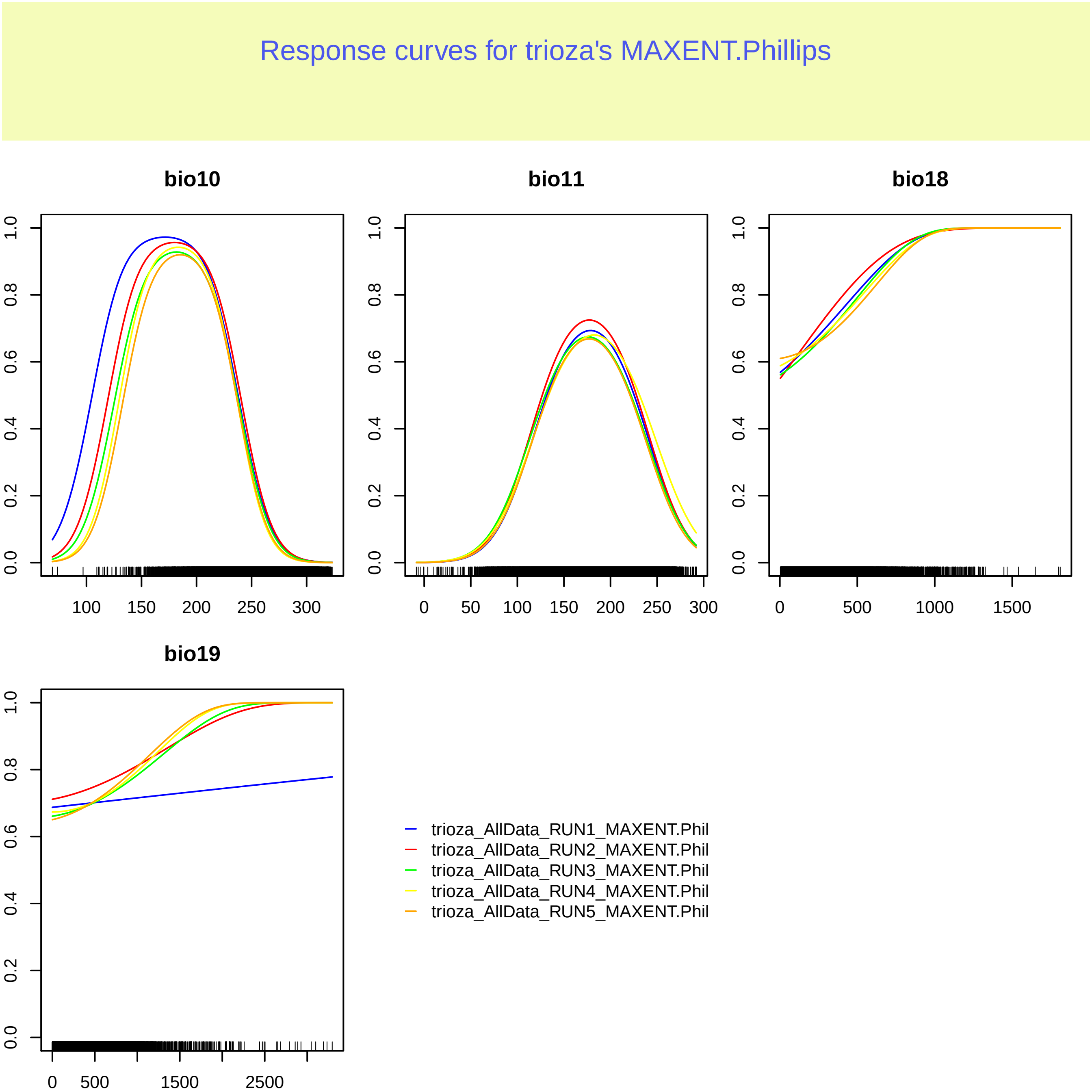

### Appendix S3

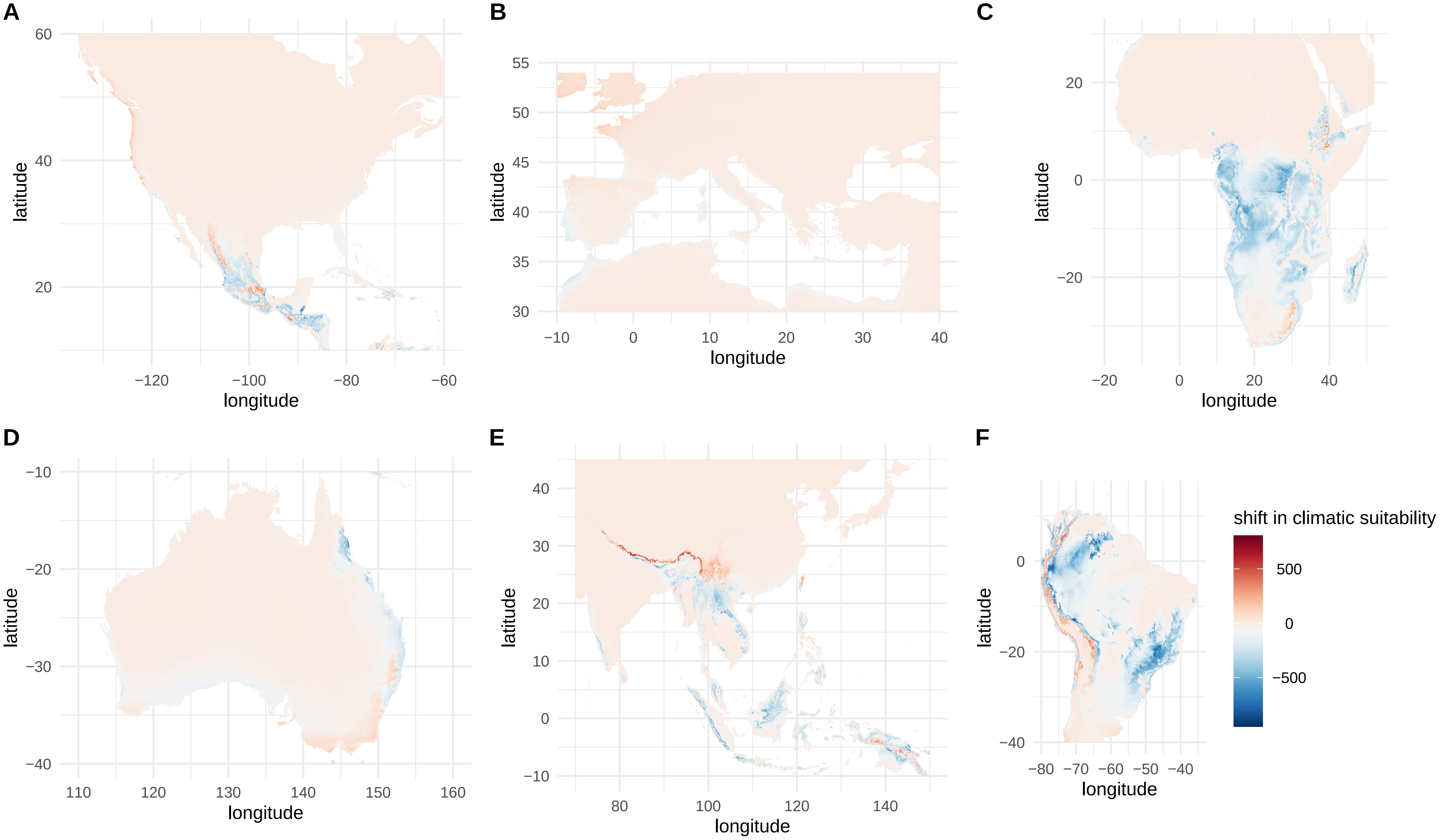
